# PKMYT1 is a computationally predicted target for kidney cancer

**DOI:** 10.1101/677245

**Authors:** Denis Torre, Nicolas F. Fernandez, Avi Ma’ayan

## Abstract

Protein Kinase Membrane Associated Tyrosine/Threonine 1 (PKMYT1) is an understudied member of the serine/threonine protein kinase family. PKMYT1 is listed as a dark kinase according to the Illuminating the Druggable Genome (IDG) target development level classification. Using a combination of bioinformatics tools that we developed, we predict that targeting PKMYT1 is potentially beneficial for treating kidney cancer.

## Introduction

While it is known that PKMYT1 is a homolog of the well-studied cell-cycle kinase Wee1, and it is involved in regulating the cell cycle^1–3^, PKMYT1 has attracted little research attention so far. Currently, as of November 2019, only 53 publications are returned when searching PKMYT1 on PubMed, compared with over 1,193 for Wee1. As a result, PKMYT1 is listed as a dark kinase according to the Illuminating the Druggable Genome (IDG) target development level (TDL) classification^4^. Here we investigate the potential role of PKMYT1 as a therapeutic target using several bioinformatics tools and databases that we developed for the IDG program.

## Results

Using the tool Geneshot^5^, which is a search engine for associating any search term with ranked lists of most relevant genes, a search for PKMYT1 returns 70 other genes, with CDK1 and WEE1 as the top two most co-occurring genes in publications that mentioned PKMYT1 as of November 2019 using the AutoRIF option. Below is a permanent link to the Geneshot search results: https://amp.pharm.mssm.edu/geneshot/index.html?searchin=PKMYT1&searchnot=&rif=autorif

Enrichr^6^ analysis of those 70 co-occurring genes from Geneshot identifies E2F4 and FOXM1 as the top transcription factors that regulate genes that co-occur in publications with PKMYT1 (ChEA-ENCODE combined library, Fisher exact test, BH adjusted p-value 5.832e-16) and G2/M transition of the cell cycle is the most enriched gene ontology (GO) Biological Process term (BH adjusted p-value 1.163e-7). Suggesting a role for this gene in the regulation of the cell cycle. Interestingly, Human Phenotype Ontology^7^ enrichment ranks nephroblastoma (HP:0002667) and embryonal renal neoplasm (HP:0011794) as the most enriched human phenotypes for these genes, suggesting that the genes related to PKMYT1 in the literature may have a more specific role in kidney related neoplasms. Below is a permanent link to the Enrichr analysis results: https://amp.pharm.mssm.edu/Enrichr/enrich?dataset=f56be87f7c63ea8f0efbff760ba6cdb6

We further confirmed this hypothesis by testing how well PKMYT1 gene expression levels, in cancer patients, can separate TCGA tumors based on their survival data (Fig. 1). We observe that PKMYT1 expression is specifically an excellent potential single biomarker to predict the outcome of kidney renal clear cell carcinoma (KIRC) based on analysis of the TCGA data. Patients with kidney carcinomas that display high expression levels of PKMYT1 also exhibit significantly poorer prognosis (log-rank test, p-value 1.92e-6, 530 samples, and 158 deaths). PKMYT1 is also highly expressed in tumors compared to normal tissue in the TCGA kidney renal clear cell carcinoma cohort, as well as in the breast, lung, and colon cancer TCGA cohorts (Fig. 2).

**Fig. 1.**
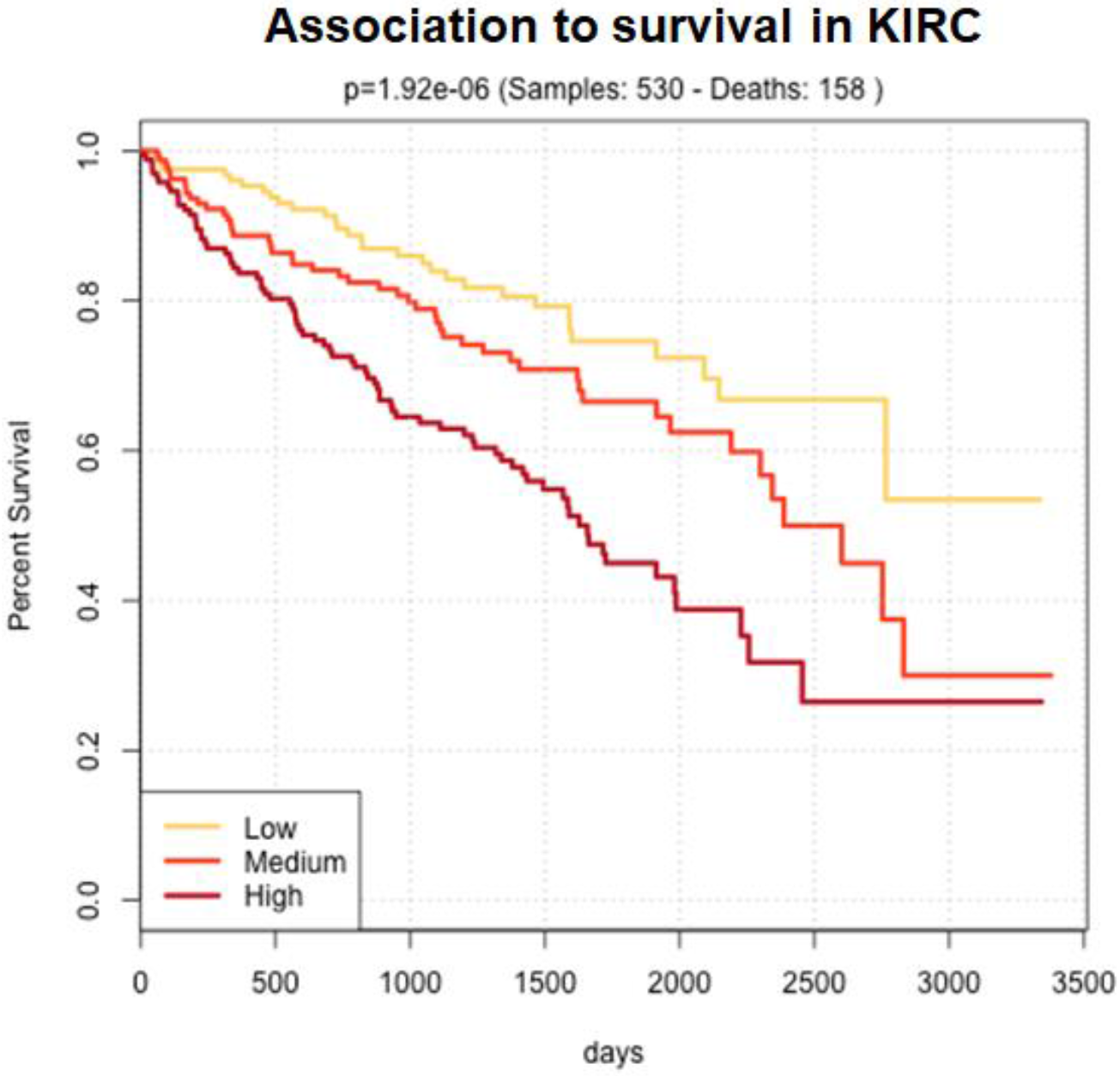
TCGA KIRC patients were divided into three even groups: PKMYT1 low, medium, and high expression to produce K-M curves. The p-value was computed with the log-rank test.

**Fig. 2.**
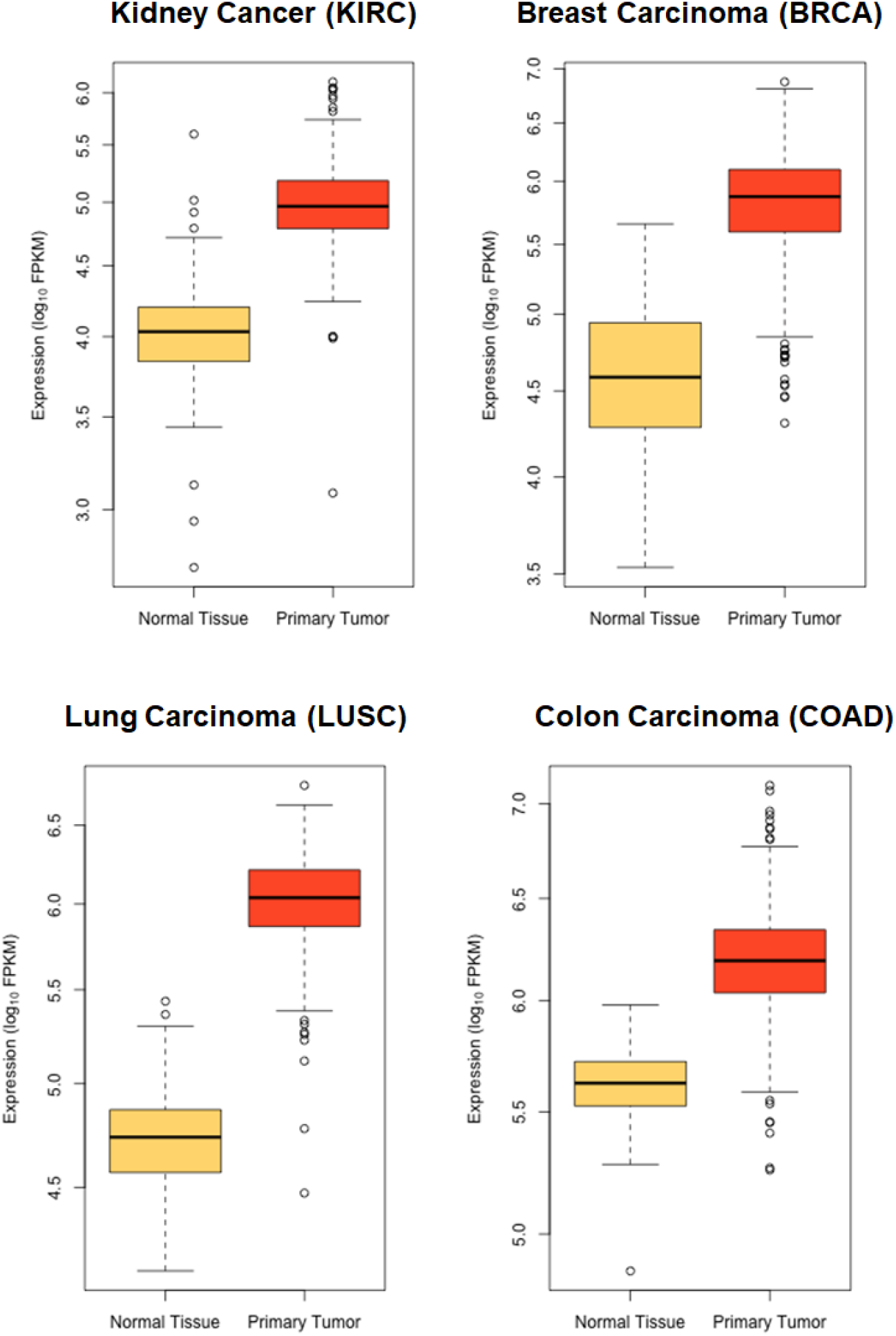
Average expression of PKMYT1 across all tumors vs. normal tissue for four cancer types from TCGA.

The involvement of PKMYT1 in the regulation of the cell cycle is further highlighted in global co-expression analysis of kinase mRNA levels across the Cancer Cell Line Encyclopedia (CCLE) dataset^8^. Such analysis identifies PKMYT1 as the only IDG dark kinase that is tightly co-expressed with a cell cycle kinome module that includes well-studied kinases such as CDK1, BUB1, AURKB, CDK2, and CDK4 (Fig. 3). Clustering analysis of the human kinome based on CCLE gene-gene co-expression data was accomplished with Clustergrammer^9^, another tool developed by the Ma’ayan Lab for the IDG-KMC. The two other dark IDG kinases found in this cell cycle module are NEK4 and PKN3, but these two dark kinases co-express less strongly with the cluster compared with the tight co-expression module that contains PKMYT1.

**Fig. 3.**
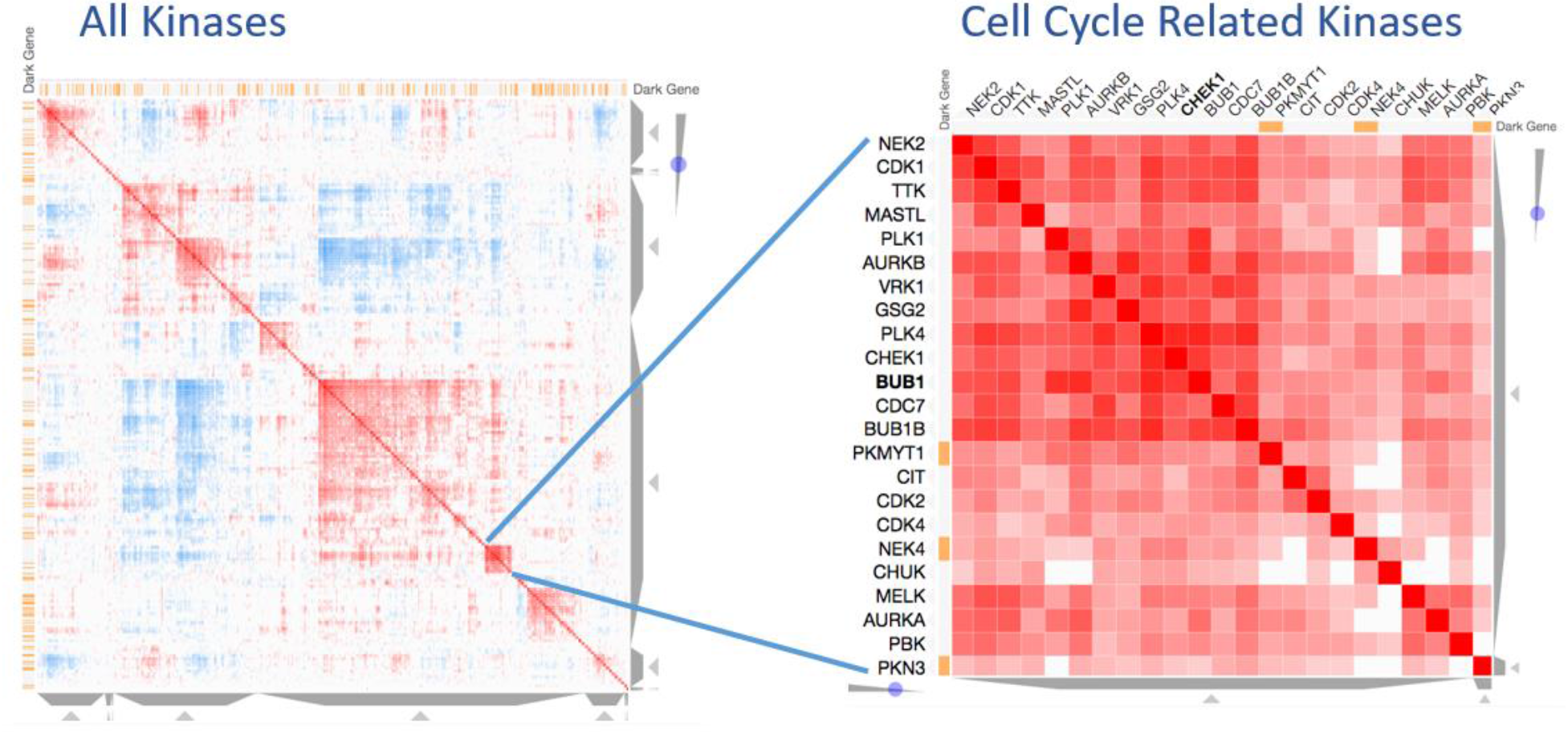
Interactive visualization of the human kinome based on mRNA co-expression correlation. Co-expression was computed with all the gene expression data from CCLE. Orange bars indicate IDG designated dark kinase.

Interactive visualizations of Figure 3 with additional plots are available from the URL below: https://maayanlab.github.io/KMC_2019/similarity_matrices.html

To identify potential small molecules that can down-regulate the expression of PKMYT1, we ran Dr. Gene Budger (DGB)^10^. DGB was developed to quickly identify drugs and small molecules that up- or down-regulate the expression of single human genes based on data collected from the original Connectivity Map^11^, the data from the LINCS L1000 assay^12^, and gene expression signature from drug perturbation followed by expression we collected to create the CREEDS resource^13^. When submitting PKMYT1 to DGB, the top drug that is suggested is pyrvinium, an FDA approved anthelmintic drug that is already being considered as a cancer therapeutic^14^. It would be interesting to test the effect of pyrvinium, and other potential novel inhibitors of PKMYT1^15^, on the proliferation of kidney cancer cell lines with the expectation of cell cycle inhibition via decrease PKMYT1 expression and activity. Along these lines, a recent CRISPR/Cas9 screen for glioblastoma identified PKMYT1 as a top hit^16^. In summary, using a combination of the IDG-KMC bioinformatics tools that we developed, we can quickly and systematically discover under-appreciated opportunities for drug and target discovery. PKMYT1 stands out as the mostly likely dark target for potential efficacy for diagnosing and treating several cancers.

## Acknowledgements

This work was partially supported by NIH grants U54-HL127624 (LINCS-DCIC), U24-CA224260 (IDG-KMC).

